# On the study of evolutionary predictability using historical reconstruction

**DOI:** 10.1101/001016

**Authors:** Sandeep Venkataram, Diamantis Sellis, Dmitri A. Petrova

## Abstract

Predicting the course of evolution is critical for solving current biomedical challenges such as cancer and the evolution of drug resistant pathogens. One approach to studying evolutionary predictability is to observe repeated, independent evolutionary trajectories of similar organisms under similar selection pressures in order to empirically characterize this adaptive fitness landscape. As this approach is infeasible for many natural systems, a number of recent studies have attempted to gain insight into the adaptive fitness landscape by testing the plausibility of different orders of appearance for a specific set of adaptive mutations in a single adaptive trajectory. While this approach is technically feasible for systems with very few available adaptive mutations, the usefulness of this approach for predicting evolution in situations with highly polygenic adaptation is unknown. It is also unclear whether the presence of stable adaptive polymorphisms can influence the predictability of evolution as measured by these methods. In this work, we simulate adaptive evolution under Fisher’s geometric model to study evolutionary predictability. Remarkably, we find that the predictability estimated by these methods are anti-correlated, and that the presence of stable adaptive polymorphisms can both qualitatively and quantitatively change the predictability of evolution.

## 1 Introduction

Predicting evolution is one of the fundamental challenges of evolutionary biology (reviewed in Lobkovsky and Koonin 2012; de Visser and Krug 2014). This question became particularly prominent with Gould’s famous thought-experiment on “replaying the tape of life” (Gould 1989), which asks how our evolutionary history would have changed if we could re-run evolution from some point in the past. Evolutionary predictability can be studied using multiple methods (de Visser and Krug 2014; Achaz *et al.* 2014), including studying populations evolving in parallel (e.g. Gould’s thought-experiment) or characterizing the likely order of mutations in a historical bout of adaptation (Weinreich *et al.* 2006) using an experimental design pioneered by Malcolm *et al.* (1990).

To study evolutionary predictability, one must be able to quantify the similarity between two different evolutionary paths. This quantification can be done in a number of different ways, which we classify using four different properties. The first property is the “type” of predictability one is studying, which can be either the predictability of the evolutionary future of a given system (“future predictability”, i.e. predictability as defined by Gould), or the predictability of the order of mutations in a completed adaptive path (“historical predictability”, i.e. predictability as inferred using the Weinreich *et al.* (2006) method). The second property is the “level” at which we are studying predictability, which can either be conducted at the genotype, phenotype or fitness level. More concretely, one can compare two evolutionary paths by the specific adaptive mutations that occurred in each path, the similarities of the organisms in some physiological trait (e.g. body-mass index), or by comparing their fitness. The third property is the “starting point” of the evolutionary paths being studied, and is only relevant for future predictability studies. In “parallelism” studies, one tries to characterize evolutionary predictability using organisms starting from similar or identical genetic, phenotypic or fitness states (e.g. when different populations of the same species independently evolve in similar ways when under similar selective pressures), while “convergence” studies characterize how organisms with distinct initial states reach similar evolutionary solutions for similar biophysical problems (e.g. the evolutionary convergence in both birds and bats to use wings for flight). The fourth property is the “resolution” of the data being used to study evolutionary predictability, which can range from a single sample of genomes from an extant population, a few major phenotypic transitions inferred from the fossil record or detailed knowledge of every single adaptive mutation that ever reached appreciable frequency in the population during the study period.

Perhaps the best known experimental study of future predictability is a set of 12 parallel *Escherichia coli* lineages that have been evolving in controlled laboratory conditions for over 60,000 generations (Wiser *et al.* 2013; Lenski *et al.* 2015; Tenaillon *et al.* 2016). In our formalism, these are studies of parallel future predictability, and analyses of this system have been conducted at the genotypic, phenotypic and fitness levels with very high resolution. One remarkable result from these experiments is that replicate independent evolution experiments frequently acquired similar adaptive mutations and experienced similar gains in fitness over the course of evolution (Crozat *et al.* 2010; Wiser *et al.* 2013), which has also been shown in other short-term laboratory evolution experiments (Tenaillon *et al.* 2012; Lang *et al.* 2013; Kvitek and Sherlock 2013; Venkataram *et al.* 2016). This suggests that evolution can be future predictable to a surprising degree over both short and long timescales.

Studying future predictability is critical to many current problems in evolutionary biology, including cancer and the development of drug resistance. However, future predictability studies are either technically or ethically infeasible in many systems of interest, leading to the development of methods to study historical predictability. While it is possible to study historical predictability at the phenotypic level (e.g. by inferring the order of phenotypic transitions from the fossil record), most recent studies characterize genotypic historical predictability. In the first major study of historical predictability, Weinreich *et al.* (2006) reconstructed all 32 possible combinations of 5 mutations in the beta-lactamase gene in *E. coli*, which are known to confer resistance to the drug rifampicin. They then quantitatively assayed the drug resistance of these 32 genotypes as a proxy for fitness, and used this information to analyze all 5! = 120 possible mutational paths (orders in which the 5 mutations could occur) from the ancestral to the resistant five-mutation genotype. A mutational path was deemed viable if resistance monotonically increased with every mutational step, and the relative likelihood of each viable path was calculated using standard population genetic methods. A number of other groups have also studied historical predictability, including studies in *Aspergillus niger* (Franke *et al.* 2011), an empirical RNA-protein binding landscape (Buenrostro *et al.* 2014), adaptive mutations identified from ancestral protein reconstruction (Bridgham *et al.* 2006; Ortlund *et al.* 2007) and adaptive mutations from a laboratory evolution experiment (Khan *et al.* 2011).

Despite these numerous studies, it is unknown whether historical predictability is truly a useful proxy for studying future predictability. While historical predictability is likely accurate in systems where adaptation is known to be limited to a very small number of mutations (e.g. the 5 mutations used by Weinreich *et al.* (2006) are the only major rifampicin resistance mutations in that system), it is unclear whether conclusions drawn from studies of historical predictability are similar to the results of studies of future predictability when not every available adaptive mutation is used for inferring historical predictability. For example, it is unknown whether the historical predictability results of Franke *et al.* (2011) in 2-6 mutation subsets of an 8 locus system are informative in understanding adaptation on the entire 8-locus fitness landscape. Due to the combinatorial nature of studying historical predictability, where 2*n* genotypes need to be considered in a system of n adaptive mutations, it is extremely challenging to study historical predictability over all known adaptive mutations in systems with highly polygenic adaptation.

In addition, all of these studies of both future and historical predictability were conducted under the assumption that all adaptive mutations successively fix in the evolving population. The presence of stable polymorphisms may significantly influence the relationship between future and historical predictability by modifying the fitness advantage of a mutation based on the alleles already present in the population. As stable polymorphisms can be generated by a variety of mechanisms, including heterozygote advantage, frequency dependent selection, and spatio-temporal variability in selective pressures, one might expect that they play a major role in the evolution of natural populations and thus influence the study of evolutionary predictability in natural systems.

In this work, we simulate adaptive evolution using Fishers geometric model (FGM, Fisher 1930; Orr 1999, 2005) to study the relationship between future and historical predictability, as well as the impact of balanced polymorphisms on the study of predictability. FGM is a phenotypic model where individuals have phenotypes defined as points in n-dimensional space. Mutations are arbitrary vectors in this n-dimensional space, allowing for an infinite number of functionally distinct possible mutations and providing an excellent model of polygenic adaptation. A fitness function is then used to map phenotypes into fitness, generating a fitness landscape that can be used to simulate adaptive evolution. FGM is a useful framework to study adaptation as it has been found to be consistent with many empirical results, including the distribution of fitness effects (DFE) of beneficial mutations, the distribution of epistasis and the effect of epistasis on the DFE, and the presence of antagonistic pleiotropy (Martin *et al.* 2007; Sousa *et al.* 2012; MacLean *et al.* 2010; Blanquart *et al.* 2014)(reviewed in Martin 2014; Tenaillon 2014). In addition, Sellis *et al.* (2011) showed that adaptive mutations in diploid FGM simulations are frequently overdominant (exhibit heterozygote advantage) if the mutations are sufficiently large in phenotype space, resulting in balanced polymorphisms. These overdominant mutations are temporarily stable, but they can be driven out of the population by subsequent adaptive mutations.

In this work, we compute both the parallel phenotypic future predictability and genotypic historical predictability of evolution from the same simulations of adapting populations to test whether future predictability and historical predictability are correlated. We then use both of these metrics to test whether overdominant mutations significantly impact the predictability of evolution. We find that these two types of predictability are anti-correlated in our simulations, and that the presence of stable polymorphisms can both quantitatively and qualitatively change our ability to predict evolution.

## 2 Model

### 2.1 Fisher’s Geometric Model Simulations

#### 2.1.1 Overview of model and simulations

In order to study predictability, we first need to generate a large number of independent adaptive trajectories. We utilize a variant of the standard haploid continuum-of-alleles Fisher’s geometric model (FGM) framework (Fisher 1930; Kimura 1965) that has been modified to consider diploid individuals (Sellis *et al.* 2011).

In FGM, we model a set of *n* independent quantitative phenotypic traits, which can be considered a vector ***r*** in ***n***-dimensional space. This vector defines an allele, thus resulting in a continuum of possible alleles across this ***n***-dimensional space. In our simulations, the initial population was defined to be monomorphic with a single fixed allele ***r_anc_***, where ||***r_anc_***|| was set to be 2 units from the optimum. Mutations in this model are defined by a mutation vector ***m***, which is used to modify an existing allele to generate a new allele. These vectors are drawn from a continuous distribution, and thus new mutations can produce an infinitely many different alleles. The phenotype vector associated with any allele can thus be calculated as ***r_anc_*** + **Σ*m_i_*** where we sum over all mutations that gave rise to the allele of interest. All mutations are assumed to be in complete linkage with each other. A haploid individual, which has one allele, has a phenotype identical to the phenotype of the allele, while the phenotype of a diploid individual is the average of the phenotype vectors of the constituent alleles. We assume sexual reproduction for diploids with free assortment of alleles and no recombination. The fitness of an individual is a spherically symmetric gaussian function of an individual’s distance from the optimum 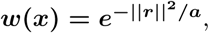 where ***a*** is a constant defined by our parameter regimes (described in the next section).

As a concrete example (Figure 1), let us consider a geometric model consisting of n = 2 phenotypic dimensions (traits 1 and 2) and considering two separate mutational events (A and B). We will begin with the haploid case (Figure 1a). The ancestral allele (anc) has a predefined phenotype (***r_ab_***, Figure 1a cross symbol) with fitness defined as a gaussian function of its distance from the optimal phenotype (||***r_anc_***||). A mutation ***m_A_*** on this ancestral allele would result in the allele A (plus symbol), defined by its phenotype ***r_A_*** = ***r_anc_*** + ***m_A_***. A further mutation ***m_B_*** on the A allele would result in the AB allele with phenotype ***r_AB_*** = ***r_A_*** + ***m_B_*** (open circle).

**Figure 1.**
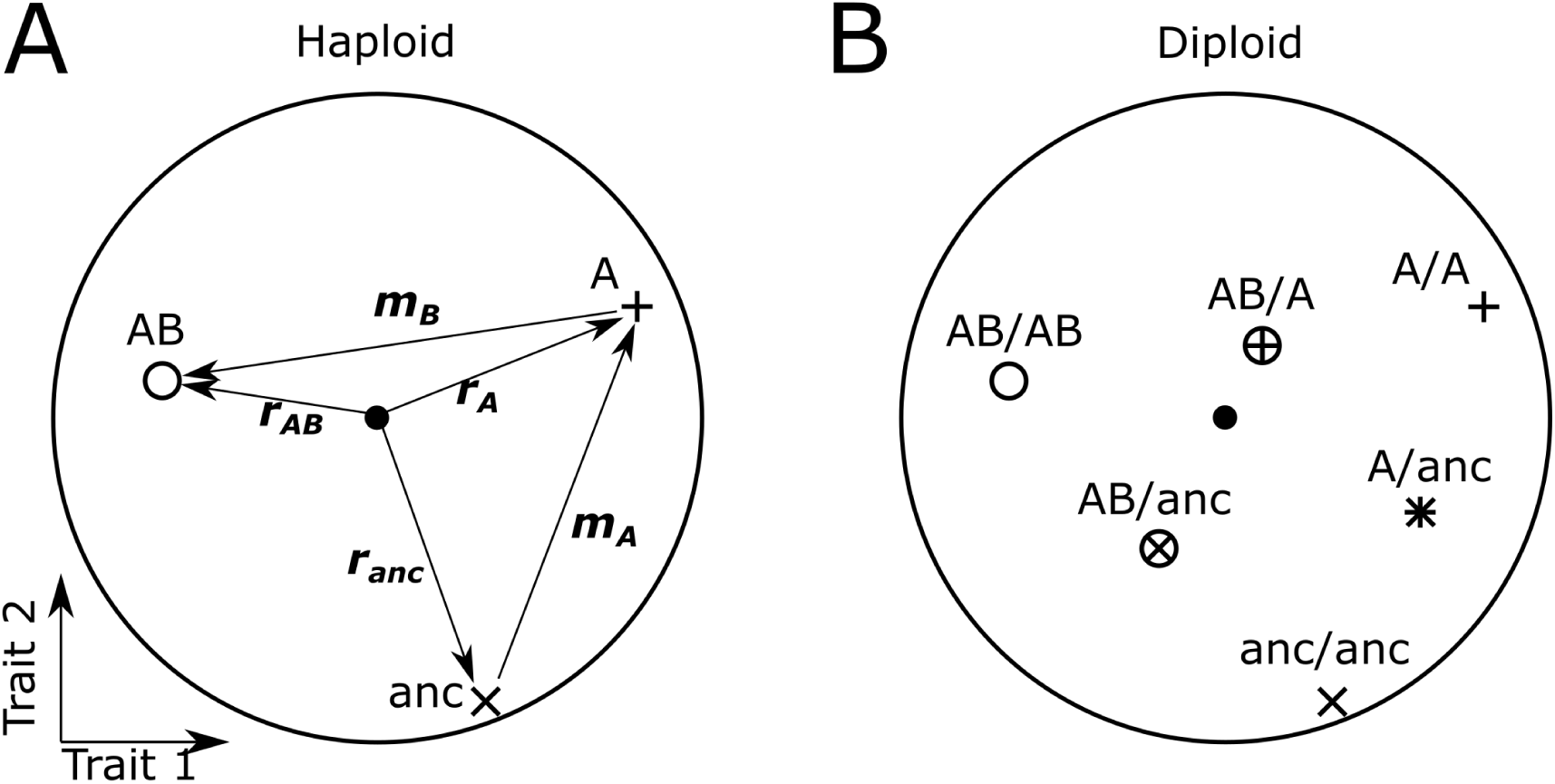
Description of Fisher’s geometric model. For illustrative purposes, we show an example of a two-dimensional Fisher’s model considering two mutations A and B. A) Haploid case. An ancestral allele, anc, is pre-defined with a set phenotype (***r_anc_***, cross), with fitness as a gaussian function of its distance ||***r_anc_***|| from the optimal phenotype (closed circle). Mutation A occurs on the background of the anc allele and adds a mutation vector ***m_A_*** to the phenotype of allele anc to get the phenotype of allele A (***r_A_***, plus sign). A further mutation B on the A allele would result in the AB allele and add the phenotypic mutation vector ***m_B_*** to the phenotype associated with the A allele and generate the new phenotype associated with the AB allele (***r_AB_***, open circle). B) We will now consider the diploid case, using the same alleles and mutation vectors as in the haploid example. For simplicty, we only show the phenotypes of the individuals as points, without the associated phenotype and mutation vectors. In this case, as individuals have multiple alleles, the phenotype of an individual with a given genotype is the average of the phenotypes of the component alleles. Thus, an individual homozygous for the A allele (A/A genotype, cross) would have the same phenotype as an A haploid individual, while an individual heterozygous for the A and anc alleles would have an intermediate phenotype (A/anc, star, the average of ***r_anc_*** and ***r_A_***).

We will now consider the diploid case (Figure 1b), using the same alleles and mutation vectors as in the haploid example. For clarity, we do not show the mutation vectors or phenotype vectors, but display the phenotypes associated with a given genotype as a point. In this case, as individuals have multiple alleles, the phenotype of an individual with a given genotype is the midpoint of the phenotypes of the component alleles. Thus, an individual homozygous for the A allele (A/A genotype) would have the same phenotype as an A haploid individual, while an individual heterozygous for the A mutant and ancestral alleles would have an intermediate phenotype (A/anc, star in Figure 1b).

This model is used to conduct forward Wright-Fisher simulations to generate the adaptive walks used for analysis throughout this work. The simulations use the code modified from Sellis *et al.* (2011) to allow for more than 2 dimensions. We perform 2,500 replicate simulations using a standard Wright-Fisher approach (Fisher 1930; Wright 1931) in both haploid and diploid populations. Haploid simulations are conducted with a population size ***N*** = **10,000**, while diploid simulations were conducted with ***N*** = **5,000**. Simulations are conducted for 10,000 generations, where each generation consists of mutating alleles and then propagating alleles to the next generation, during which we also impose selection. Individuals to be mutated are uniformly sampled according to the mutation rate ***µ*** = **5 ∗ 10^−6^**, while the mutation vectors ***m*** have a magnitude drawn from the exponential distribution with ***λ*** = **2** and a uniformly distributed direction. Propagation is conducted via sexual reproduction in diploids with the possibility of selfing, with an implicit assumption of random mating between all individuals present in the current generation to compute the genotype frequencies of the offspring. We assume that there are an infinite number of offspring, which are then multinomially sampled in a manner weighted by both the frequency of each genotype in the offspring pool and the fitness of that genotype to give rise to the population present in the next generation (i.e., viability selection on the offspring). Note that this sexual reproduction process is only meaningful in the diploid model to compute the offspring pool genotype frequencies. In haploids, the genotype weights used in the multinomial sampling pool are identical to their frequencies in the current generation multiplied by their fitness.

For all of our statistical analyses, we considered only those mutations that are present on the most frequent allele at generation 10,000. These mutations are the set of mutations present in the adaptive walk that are most easily sampled in natural populations. The low per-generation mutation rate (0.05 mutations per generation) allowed us to use the strong-selection weak-mutation (SSWM) assumption for our analysis to consider each mutation by itself. This assumption is consistent with biological systems with small effective population sizes which results in periodic fixation of beneficial mutations. Some examples of systems with small enough effective population sizes to be in this regime (tens of thousands of individuals or less) include humans (Tenesa *et al.* 2007), microbial communities in nectar flowers (Herrera *et al.* 2009) and many species listed by the IUCN as endangered or vulnerable (IUCN 2016). We additionally limited our analysis to the first five mutations of each adaptive walk in order to compare adaptive walks of equal lengths and re-ran simulations with fewer than five mutations until they met this criteria. This is comparable to many recent studies studying genotypic historical predictability (Weinreich *et al.* 2006; Khan *et al.* 2011; Franke *et al.* 2011). For simplicity of analysis, diploid simulations that generated stable polymorphisms containing three or more alleles were discarded and re-run until they met this criteria. We partitioned the diploid simulations into those that did and did not contain overdominant mutations to study the impact of stable polymorphisms on the predictability of evolution (described in a subsequent section).

#### 2.1.2 Parameter regimes

In all of our simulations, the population initially contains a single allele with a distance of 2 units from the optimum. Since we are using only spherically symmetric fitness functions, the exact position is irrelevant. We conduct our first set of simulations in a two dimensional regime (n = 2) with a poorly adapted initial population that is far from the optimum (2D-Far regime). The gaussian fitness function for this regime is defined with ***a*** = **2**. We conduct additional simulations in two additional regimes to validate our results: one regime where the population is initialized close to the phenotypic optimum (2D-Close regime, ***a*** = **18**), and one regime where the population is evolving in 10-dimensional space rather than 2-dimensional space (10D-Far regime). The 2D-Close regime was selected such that the initial population, at 2 units from the optimum, is in the “concave-down” portion of the fitness surface.

#### 2.1.3 Partitioning adaptive walks

In order to explore the effect of overdominance on predictability we have separated the diploid simulations into those with and without overdominant mutations. The methodology for this separation is based on the fact that all overdominant mutations must be capable of creating a stable polymorphism with two alleles, so by inferring whether or not a mutation can create such a stable polymorphism, we can infer whether or not it is overdominant. We begin with the five-mutation adaptive walks identified previously and first determine the time ***t*_5_** at which the allele containing these first five mutations exceeded **5%** frequency in the population. All time-points after ***t*_5_** are no longer considered for analysis. At each generation ***t*** ≤ ***t*_5_**, we isolate all alleles in the population at **≥ 1%** frequency. For every subset of these alleles, we compute their equilibrium frequencies and mean fitness using the method of Kimura (1956).

Briefly, this is done by computing an n x n matrix A for the n alleles under consideration, where the value of ***A_ij_*** is the fitness of the genotype defined by alleles i and j. We also consider a matrix T, where ***T_ij_*** = ***A_ij_*** − ***A_in_*** − ***A_jn_*** + ***A_nn_***. If we denote the equilibrium frequency of the ith allele by ***x_i_***, we get 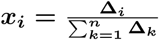 where **Δ_*i*_**, is the determinant made by substituting ones for all the elements of the ith column in the matrix A. With these equilibrium frequencies, we can easily calculate the mean fitness at this equilibrium as 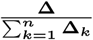 where **Δ** is the determinant of A.

The necessary and sufficient conditions for determining whether a set of n alleles can make a stable equilibrium in the first place are: 1) that the quadratic form T be negative definite and 2) **(−1)^*n*−1^Δ_*i*_** > **0** for all i = 1, 2, .. , n. For further reading, please see Kojima (1959); Mandel (1959); Kingman (1961).

If a set of alleles generates a stable polymorphic state at equilibrium, we infer that there is an overdominant mutation present among those alleles. An FGM simulation is determined to contain an overdominant mutation if, for any generation ***t*** ≤ ***t***_**5**_, the subset of alleles with the highest mean fitness at generation t is a stable polymorphism at equilibrium. For simplicity, we removed simulations that contained stable polymorphisms with ≥**3** alleles for ≥**50** generations so that we only need to consider 2 allele stable polymorphisms for the remainder of this work.

For the diploid simulations in each parameter regime, we ensured that at least 500 of the simulations did not contain any overdominant mutations (the identification of which is described in the next section) by simply rerunning some of the simulations until this criteria was met. This was done to ensure that we had a sufficient number of simulations with and without overdominant mutations for statistical analysis.

The rationale for this approach is as follows. An allele generated by an overdominant mutation that successfully invades the population must produce a stable polymorphism. This is also the only means by which a stable polymorphism can be generated in our simulations, as we do not allow for any other mode of balancing selection. However, it is not possible to robustly detect stable polymorphisms simply by tracking the frequencies of the mutations in the population and detecting whether they are being maintained in the population, due to (rare) issues with clonal interference (Desai and Fisher 2007; Herron and Doebeli 2013; Kvitek and Sherlock 2013; Lang *et al.* 2013). It is also not possible to directly infer that an allele is overdominant by simply comparing the fitness of different genotypes to detect heterozygote advantage as there are potentially more than two alleles present in the population when the mutation reaches substantial frequency either due to clonal interference or due to a mutation invading the population when there is already a stable polymorphism from a prior overdominant mutation.

Therefore, we need to separately test whether each new mutation in the population could result in a stable polymorphism if no additional mutations were allowed (eliminating the clonal interference problem). Since there are an arbitrary number of alleles present in the population at any one time (again, due to clonal interference), we make a simplifying assumption that the set of alleles will not result in an equilibrium resulting in more than 2 alleles being stably maintained in the population, which is valid as we have prescreened all of our simulations to eliminate any that contain stable polymorphisms of more than 2 alleles.

We can now utilize the Kimura method for calculating both whether a set of alleles could be stably maintained at equilibrium and the mean fitness of the population at that equilibrium for determining 1) whether the new mutation can invade the population and 2) whether it will be maintained as a stable polymorphism if it does invade. The set of alleles under consideration is all of the alleles that already existed in the population and the new allele generated by the new mutation. We use the Kimura method on all pairs of these alleles and identify the pair that generates the highest mean fitness at equilibrium. This highest fitness equilibrium state is then used to determine whether 1) the new allele successfully invaded (if the pair with the highest equilibrium fitness results in an equilibrium state that does not include the new allele, it cannot invade the population) and 2) whether it is being maintained as a stable polymorphism or has fixed in the population (the presence of a stable polymorphism implies that the mutation that generated the new allele was overdominant).

#### 2.1.4 Identification of hidden alleles

For each generation of every FGM simulation, we identified the expected equilibrium state of the population considering only those alleles at >**1%** frequency at that generation in section 2.1.3. We then identify the hidden alleles for a simulation as alleles in the equilibrium population states for all generations ***t*** < ***t***_**5**_ with that does not solely consist of a subset of the five mutations under consideration. In other words, hidden alleles are those alleles that reach substantial frequency before ***t*_5_** and would have been present in a stable equilibrium if they had not been out-competed by a later allele and were thus excluded from our original identification of the 5-mutation adaptive walk.

### 2.2 Studying future predictability

We first quantify the average adaptive walk in phenotype space for each ploidy and parameter regime used in our simulations. Given that all of our adaptive walks consist of exactly five mutations, the average adaptive walk also consists of five mutations, where the first mutation vector is the average of all of the observed first mutation vectors across all simulations, the second mutation vector in the average walk is the average of all observed second mutation vectors, and so on. This average adaptive walk matches the expected adaptive walk of a straight line in phenotype space leading from the ancestral phenotype directly towards the fitness optimum. We use the average instead of the straight line as the average is directly measurable from experimental data while the line requires comprehensive knowledge of the fitness landscape, the knowledge of which would eliminate any need to study the predictability of the system.

In order to compute a summary statistic for the phenotypic parallel future predictability of an adaptive walk, we calculate the minimum distance of each observed allele during the adaptive walk from the average walk, and then taking the maximum across all observed alleles of these minimum distances as a measure of the deviation of the adaptive walk from the average walk in phenotype space. An adaptive walk that has a smaller deviation is more predictable than one that has a larger deviation.

### 2.3 Studying genotypic historical predictability

Studies of genotypic historical predictability seek to reconstruct the order in which a set of mutations arose in an adaptive trajectory. We can then study the probability distribution of all of the possible adaptive trajectories to understand how predictable the system is overall.

#### 2.3.1 Computing the likelihood of a particular order of mutations

We begin with an overview of the historical predictability method used by Weinreich *et al.* (2006) for inference when assuming that each adaptive mutation fixes in the population, and then continue on to a description of our implementation of the method which is suitable when stable polymorphisms are possible. As mentioned before, we expect stable polymorphisms to frequently occur in our diploid simulations. We explicitly model the phenotypes of the alleles and mutation vectors, and use the same fitness functions as in the FGM simulations to compute fitness.

##### Weinreich *et al* (2006) inference method

Weinreich *et al.* (2006) describe the probability of the ancestral allele (***A^wt^***) evolving into the derived allele containing all 5 mutations available (***A^der^***) going through a particular order of mutations (***M_i_***) with intermediate alleles ***a***, ***b***, ***c*** and ***d***. This can be computed as

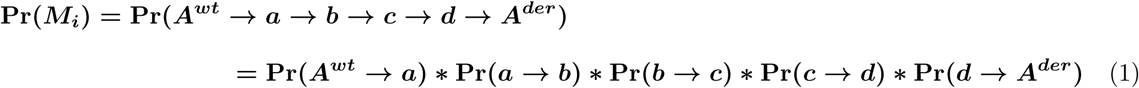

because “along any particular trajectory the choice of each next fixation is statistically independent of all previous fixations. Here, the **Pr(*i* → *j*)** are the conditioned fixation probabilities of a particular single mutant neighbor ***j*** of an allele ***i*** given by

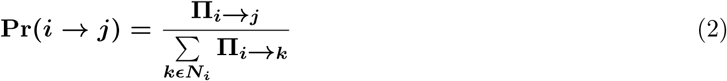

where **Π**_***i* → *j***_ is the unconditioned fixation probability of allele ***j*** from allele ***i***, and ***N_i_*** is the set of all mutational neighbors of allele ***i***.” (modified from Weinreich *et al.* (2006) Supplementary Methods). In essence, (Weinreich *et al.* 2006) compute the probability of a particular order of mutations as the product of the probabilities of each mutation in that order successfully fixing in the population in succession.

##### Our method to study genotypic historical predictability

As our simulations violate some of the assumptions of the historical predictability inference method of Weinreich *et al.* (2006), we need to modify the method to account for these violations (please see the supplementary text for a detailed example of the implementation described in this section). First, since we are using a diploid model, new mutations occur as heterozygotes and thus must invade the population as heterozygotes. Therefore, we cannot compute the fixation probability, but must compute the probability of an allele successfully invading the population from low frequency and reaching its equilibrium frequency. Secondly, in the presence of a stable polymorphism, new mutations can occur on multiple available backgrounds. This allows for the generation of hidden alleles. This also implies that it may take more than 5 mutations in a mutation order to generate the allele with all 5 mutations. Finally, a new mutation that successfully invades can either fix or balance with any of the alleles already present in the population, violating the Weinreich *et al.* (2006) assumption that the fitness of an allele is independent of the other alleles already present in the population. Therefore, we cannot treat the adaptive walk as a series of independent steps but need to take an integrated approach to study historical predictability. As it is challenging to describe the method using closed form analytic equations as in Weinreich *et al.* (2006), we will describe the recursive algorithm we use to implement the method using pseudocode. Every call to the algorithm requires a population state (set of alleles and their frequencies), a set of alleles observed during the recursion and the probability of the mutation order so far. Using global variables outside of the algorithm, we keep track of **Φ(*M_i_*)**, the unconditioned probability of every possible mutation order ***M_i_***. All **Φ(*M_i_*)** are initialized to **0**.

#### Historical predictability inference(***S_existing_*, *A^existing^*,*P_existing_***)

~~~
1: ***S_existing_*** ← the population state = a set of alleles and their frequencies
2: ***A_existing_*** ← the set of alleles observed so far in this mutation order
3: ***P_existing_*** ← the unconditioned probability of this order of mutations so far
4: if ***A^der^*** ε ***S_existing_* then**
5: We need to first determine the order ***M_i_*** in which the mutations were introduced into ***A^der^*** and add
   ***P_existing_*** to the unconditioned probability for this order of mutations (**Φ(*M_i_*)**)
6: return // We are done since we have successfully generated ***A^der^***
7: **else**
8: *ρ_total_* = 0
9: **for all** new alleles ***A^n^*** that can be generated by a single mutation on the alleles in ***S_existing_***, excluding those where ***A^n^*** ∈ ***A_existing_* do**
10: **for all** pairs of alleles ***A^i^***, ***A^j^*** in the set of alleles including ***A^n^*** and every allele in ***S_existing_* do**
11: Compute the frequency of ***A^i^*** and ***A^j^*** and the mean fitness of the population at equilibrium assuming these are the only two alleles in the population
12: ***S_new_*** = the pair of alleles and their frequencies with the highest mean fitness computed in the preceding for loop excluding all alleles at frequency 0
13: **if *A^n^*** ∉ ***S_new_* then**
14:    ***A^n^*** cannot invade ***S_existing_*** and can thus be ignored
15: **else**
16: compute 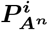 = the probability of invasion of ***A^n^*** into ***S_existing_*** through 10,000 forward Wright-Fisher simulations
17: The unconditioned probability of ***A^n^*** succeeding in this population ***ρ_n_*** = 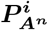 * the frequency of the allele in ***S_existing_*** that was mutated to generate ***A^n^***
18: *ρ_total_*+ = ***ρ_n_***
19: **for all** new alleles ***A^n^*** with ***ρ_n_* > 0 do**
20: ***S_new_*** and ***ρ_n_*** defined as above for ***A^n^***
21: **if** using the Weinreich et al method **then**
22: ***S_new_* = *A^n^*** at frequency 1 (fixation)
23: ***A_new_*** = ***A_existing_*** ∪ ***A^n^***
24: 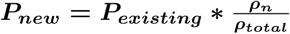
25: Historical predictability inference(***S_new_***, ***A_new_***, ***P_new_)*** // recursive call
~~~

The initial call to this algorithm has ***S_existing_*** be the ancestral population used in the FGM simulations i.e. a population monomorphic for an allele two units from the optimum, ***A_existing_*** is the set containing the single element ***A^wt^*** and ***P_existing_* = 1**. Once we have computed the unconditioned probability **Φ(*M_i_*)** for every ***M_i_***, we then use this information to compute the conditioned probability for each mutation order.

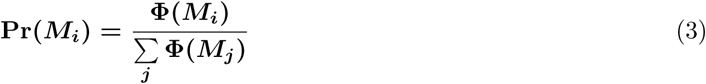

Note that we track mutation orders by the order in which the mutations were introduced on allele ***A^der^***, which is always five mutations long, not the order in which the mutations were introduced in the population in the algorithm which is >= **5** mutations. If multiple recursions through the algorithm use the same order of mutations in ***A^der^***, the likelihoods from all of these recursions are summed to get **Φ(*M_i_*)** for that particular ***M_i_***.

We compute the invasion probability of a new allele 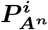 using 10,000 forward Wright-Fisher simulations. In these simulations, we set N = 5,000 diploid individuals as in our FGM simulations, with no new mutations allowed.

The probability of a new allele successfully invading and reaching the deterministically inferred stable equilibrium is then the fraction of Wright-Fisher simulations where ***A^n^*** reaches **90%** of its expected equilibrium frequency in ***S_new_***. These simulations are entirely separate from the FGM simulations used to generate the adaptive walks used throughout the rest of this work. We are forced to utilize empirical estimations through simulations and not the classical analytic solutions to compute 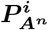 (Haldane 1927; KIMURA 1962) as many of the observed mutations have a selective advantage exceeding **100%**, violating the assumptions of the analytic solutions that the mutations are weakly beneficial. Our simulations suggest that the analytic solutions significantly overestimate the invasion probability under these conditions (data not shown).

We study historical predictability in our haploid simulations using a similar algorithm to that used for the diploid simulations. The major differences are that we 1) consider only single alleles instead of pairs of alleles when identifying the equilibrium population after a mutation and 2) empirically compute the invasion probability of a new mutation using forward Wright-Fisher simulations using a population of N = 10,000 haploid individuals instead of using a diploid model.

##### 2.3.2 Quantifying genotypic historical predictability

To quantitatively study the results of our genotypic historical predictability analysis across simulations, we define the effective number of paths statistic as

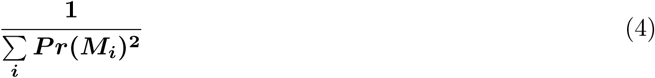

The effective number of trajectories is defined to be 0 when there are no viable trajectories, i.e. when 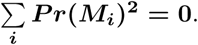 This is similar to the effective number of alleles in a population (Kimura and Crow 1964), the predictability metric of Roy (2009) and the entropy metric of Palmer *et al.* (2013).

When a single trajectory dominates the probability density, the effective number of trajectories is close to 1, indicating high historical predictability. On the other hand, if every trajectory has equal probability, 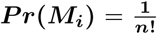 since we know that there must be ***n*!** possible mutation orders for a system of ***n*** mutations. In this situation, the effective number of paths = ***n*!** = total number of possible mutation orders, indicating low historical predictability. This provides a single metric of the diversity of mutational orders that are possible while accounting for their relative likelihoods and summarizes the historical predictability of the adaptive walk.

#### 2.4 Source Code

Complete source code for the FGM simulations is available at https://github.com/sunthedeep/Fisher-Geometric-Model.

### 3 Results

In this work, we study the predictability of evolution when adaptation can be highly polygenic, such that comprehensively sampling the entire fitness landscape is combinatorially infeasible. We compute the phenotypic parallel future predictability of an FGM simulation by comparing it to the average simulation from the same parameter regime (see Model section 2.2 for details). Simulations with a larger deviation from the average are thus less future predictable. We then compare these results to our effective number of paths statistic, which measures genotypic historical predictability. This statistic captures the distribution of the likelihoods of all possible orders of mutations observed in a given simulation (see Model section 2.3.2 for details). If measuring historical predictability is useful in inferring the future predictability of evolution, then the historical predictability of a simulation should be positively correlated with its future predictability.

#### 3.1 Comparison of future and historical predictability

We first correlate future and historical predictability using both haploid and diploid simulations in three different parameter regimes (2-dimensional regimes close and far from the optimum, and a 10-dimensional regime far from the optimum, see Model section 2.1.2 for details). We find a strong and significant negative correlation in all of these comparisons (Figure 2). In other words, adaptive walks that are highly historically predictable are future unpredictable as they are highly divergent from the average adaptive trajectory.

**Figure 2.**
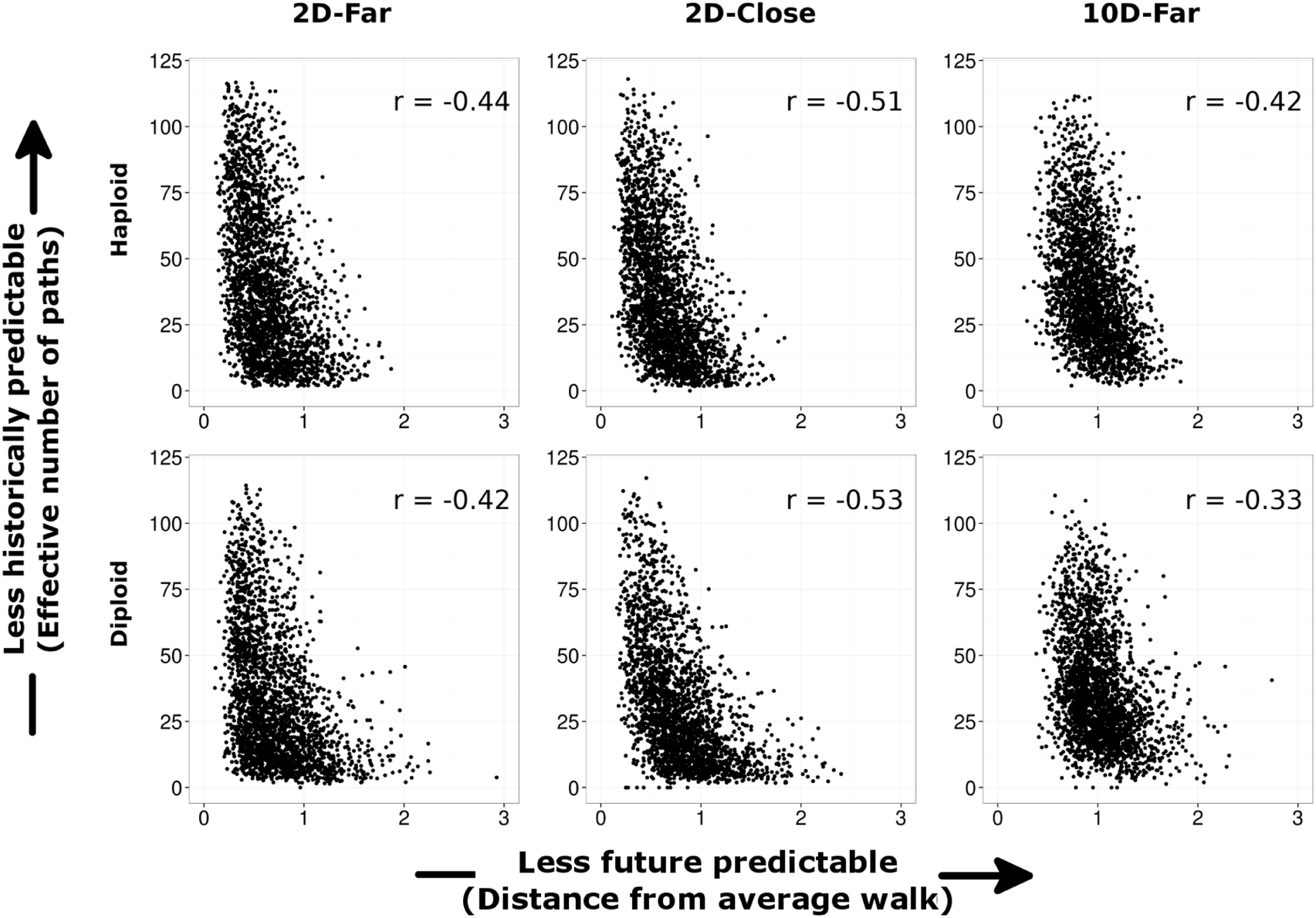
Comparison of historical and future predictability. We plot future predictability, as measured by the distance from the average trajectory metric, and historical predictability, as measured by the effective number of paths metric, for all of our simulations. We perform these comparisons across simulations from two different ploidies and three different parameter regimes. In all of these simulations, the two types of predictability are significantly anti-correlated (***p*** < **10^−10^** in all cases).

#### 3.2 Stable polymorphisms and evolutionary predictability

We also study the impact of stable adaptive polymorphisms on the predictability of evolution. In our FGM simulations, stable polymorphisms are generated through overdominant mutations in our diploid simulations. In each of our three sets of diploid simulations, we separate the adaptive walks into those that do and those that do not contain a stable polymorphism using the method of (Kimura 1956) (see Model section 2.1.3 for details). We can then compare the distributions of our historical and future predictability metrics between these two groups to test for a significant effect. We find that simulations with stable polymorphisms are significantly less future predictable than simulations without such polymorphisms (Figure 3, top row), which is concordant with the results of Sellis *et al.* (2011) that overdominant mutations allow an adaptive walk to explore a larger portion of the fitness landscape. In contrast, we find that simulations with overdominant mutations are significantly more historically predictable than simulations without such mutations (Figure 2, bottom row). These results again show the anti-correlation between future and historical predictability, and suggest that stable polymorphisms significantly impact the predictability of evolution.

**Figure 3.**
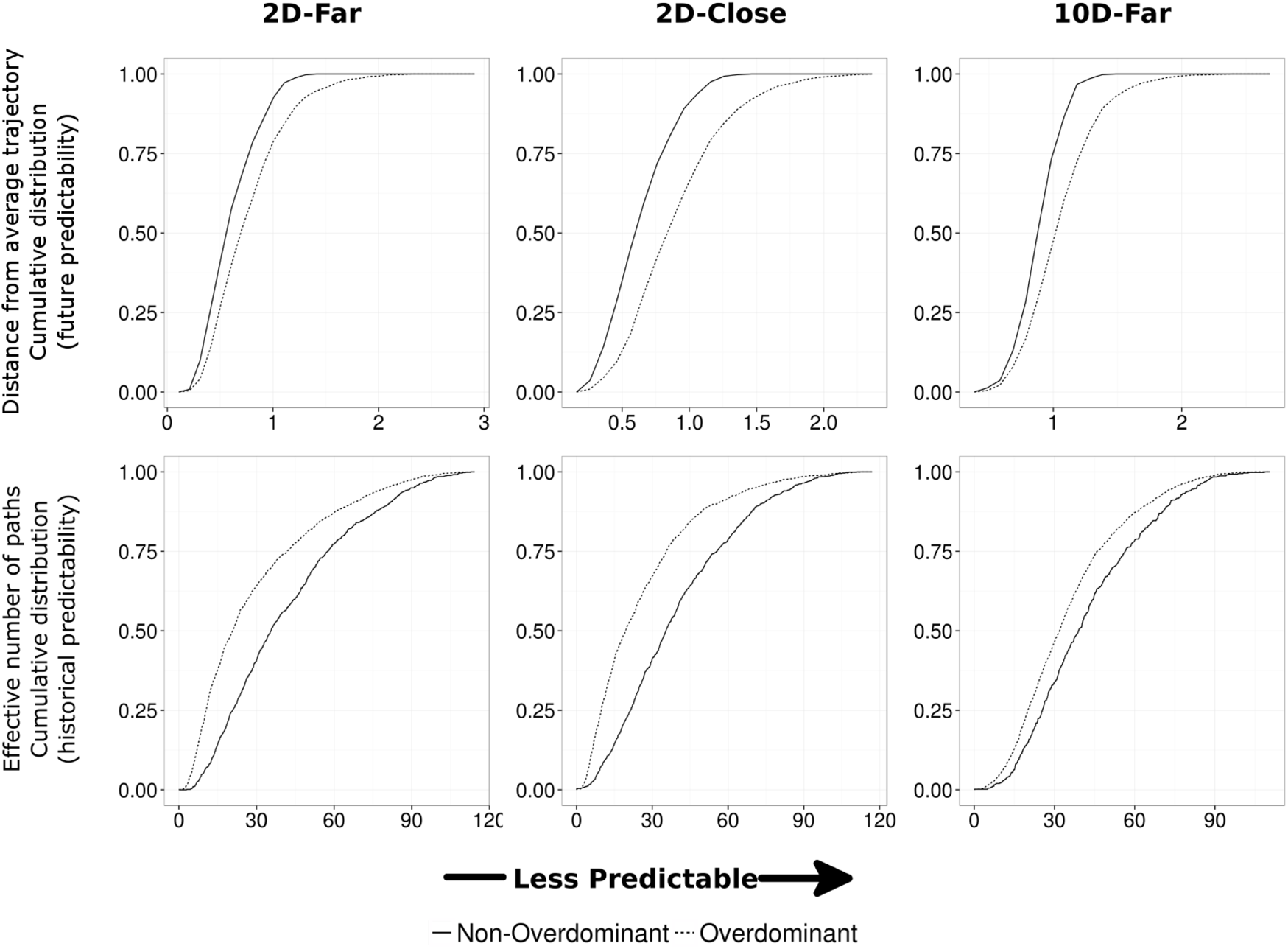
Impact of overdominant mutations on predictability. We plot the cumulative distribution of our measures of future predictability (top row) and historical predictability (bottom row) across all three diploid parameter regimes where we separate simulations that do and do not contain overdominant mutations (dotted and solid lines). We see that overdominant mutations cause adaptive walks to be significantly less future predictable but more historically predictable than simulations without overdominant mutations (Kolmogorov-Smirnov test ***p*** < **10^−10^** in all cases).

#### 3.3 Historical contingency generated by stable polymorphisms

In the course of our analysis, we were struck by the presence of 70 diploid simulations across all three parameter regimes where historical predictability analysis showed that the order of mutations that actually occurred in the FGM simulation was inviable. While some of these instances are due to multiple mutations, i.e. a population gaining a mildly deleterious mutation and then quickly gaining a highly beneficial mutation on the same background, the remaining simulations contain derived alleles that reached high frequency through a stable polymorphism but were eventually lost, which we term “hidden” alleles (Figure 4a). We hypothesized that in these simulations, hidden alleles were necessary to the evolutionary path, and not considering these alleles leads to the mistaken inference that no order of mutations was viable (Figure 4b). To test this, we selected one of these simulations at random to recompute its historical predictability while including all hidden alleles in the inference. This modification allowed us to successfully infer that the order of mutations observed in that FGM simulation was viable (data not shown), suggesting that our inability to detect hidden alleles in most systems can lead to significant errors in inference. In general we find that ≥ **25%** of diploid simulations in each of the three parameter regimes contain at least one hidden allele (see Model section 2.1.4 for details). While we find no evidence for hidden alleles generating a systematic effect on historical predictability in our model, the finding that some rare bouts of evolution are highly dependent on detecting hidden alleles highlights their potential impact in natural systems.

**Figure 4.**
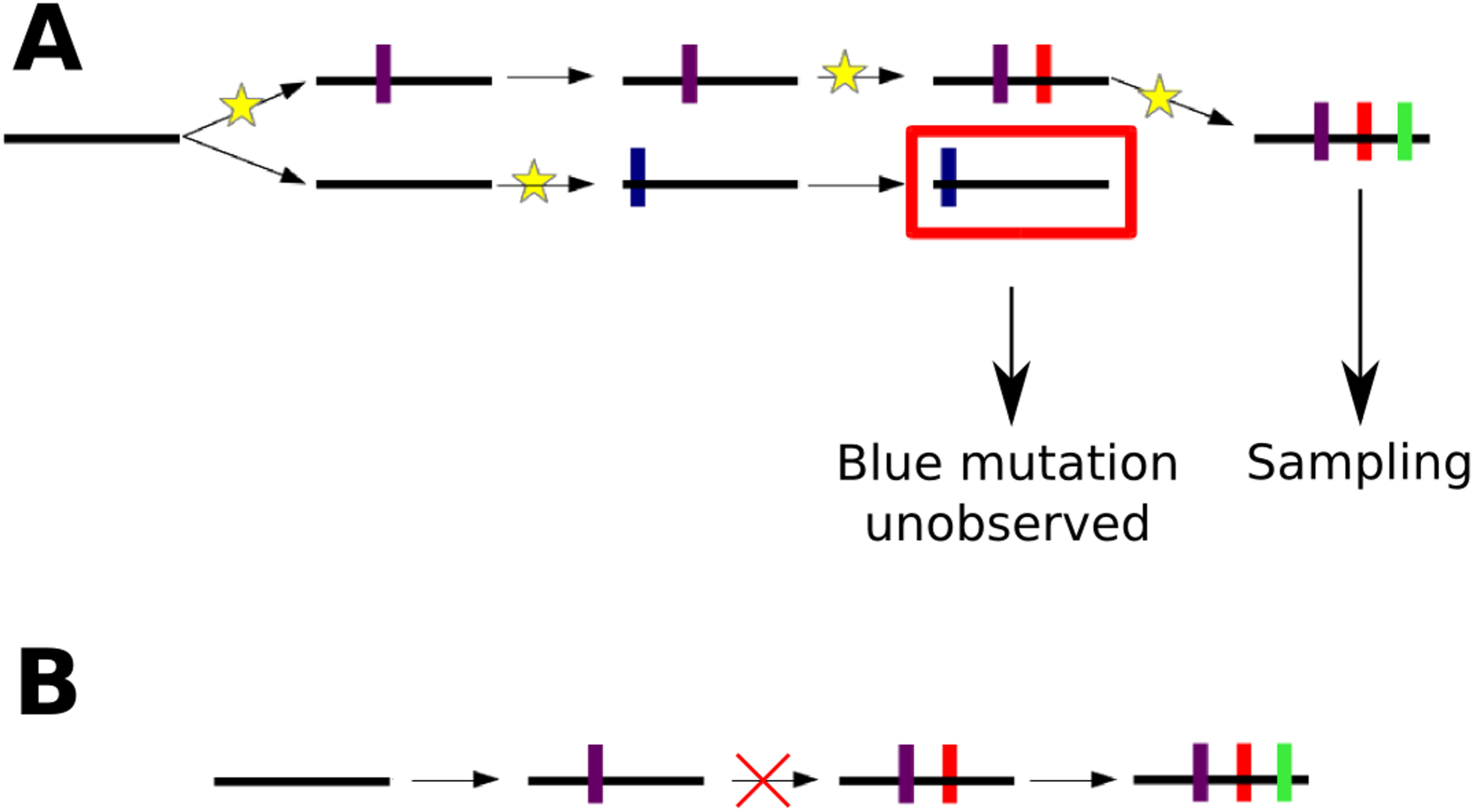
Qualitative impact of polymorphism on adaptive trajectories. Black horizontal lines represent alleles, while colored vertical bars represent different mutations. Yellow stars show the occurrence of mutations. Time increases from left to right, and each set of arrows represents a transition from one population state to another through the process of a mutation successfully invading the population and reaching equilibrium. **(A) Hidden Alleles.** A four mutation system is depicted, where the blue mutation creates a derived allele that occurred on a polymorphic state that was stably maintained for some time but subsequently lost. As this allele contains a mutation not present in the sampled allele, we call the allele with the blue mutation a hidden allele, as it is hidden from sampling. **(B) Impact of hidden alleles** In some cases, the blue allele may have been necessary for the purple, red and green mutations to occur in the order that they did, resulting in an adaptive trajectory that is impossible to accurately reconstruct without knowledge of the hidden allele.

### 4 Discussion

In this work, we sought to answer two major questions when studying the predictability of evolution in highly polygenic systems. First, we wanted to study the relationship between future and historical predictability. Second, we wanted to investigate how predictability changes when comparing simulations with and without stable polymorphisms.

In our simulations, we found that future and historical predictability are anti-correlated. This anti-correlation can be intuitively understood in the FGM framework used to conduct our simulations. Adaptive walks that are phenotypically similar to the average walk (high future predictability) tend to move relatively directly from the ancestral phenotype to the optimal phenotype on the fitness landscape. As each mutation in the adaptive walk changes the phenotype of the individual in a similar direction, there is very little sign epistasis between these mutations. A recent study has also shown that the amount of sign epistasis is correlated with the distance of the population from the optimal phenotype (Blanquart *et al.* 2014), consistent with our observations from the 2D-close and 2D-far parameter regimes (Figure S3). As previous work has shown that historical predictability is highly correlated with the amount of sign epistasis present in a system (Weinreich *et al.* 2005), high future predictability results in low historical predictability (i.e. most orders of these mutations are viable and have similar likelihood). In contrast, adaptive walks that are highly divergent from the average walk (low future predictability) are more likely to have mutations with sign epistasis that constrain their order, resulting in high historical predictability.

We can gain a similar intuitive understanding of the effect of overdominant mutations on predictability in our model. Overdominant mutations tend to overshoot the phenotypic optimum (Sellis *et al.* 2011), resulting in low future predictability and requiring subsequent mutations to be compensatory (i.e. move the phenotype in the opposite direction of the overdominant mutation). This generates large amounts of sign epistasis in simulations with overdominant mutations, resulting in high historical predictability compared to adaptive walks without overdominant mutations.

While previous work has shown how underdominant mutations in a metapopulation can generate priority effects (Altrock *et al.* 2010), our work is the first to show how overdominant mutations can generate historical dependencies in the evolutionary trajectory. The presence of transient stable polymorphisms during adaptive evolution create unsampled “hidden” alleles, which, in extreme cases, makes us infer that the true order in which the mutations occurred in the adaptive walk is inviable. This presents significant problems when attempting to study historical predictability in natural systems with a single time-point sampling resolution, as our inability to detect hidden alleles in such systems would make any inferred likelihood suspect. Studies of historical predictability under such conditions would allow the calculation of mutation order likelihoods for the set of mutations under consideration, however, we would have no idea whether these likelihoods are accurate, nor would we have any estimate of how inaccurate they are due to the missing “hidden” alleles. Functionally important hidden alleles are challenging to identify even in extant populations, due to the vast amounts of variation present in any natural population, making this a particularly difficult problem to solve.

Our simulations focused on one mechanism for the generation of stable adaptive polymorphisms, namely heterozygote advantage, but a number of other mechanisms exist in nature. These include negative frequency dependent selection (Levin *et al.* 1988; Iserbyt *et al.* 2013), and spatially or temporally variable selection (Rainey and Travisano 1998; Kasumovic *et al.* 2008; Saltz and Nuzhdin 2014). Natural populations can also generate unstable polymorphisms through a number of other mechanisms, such as clonal interference (Desai and Fisher 2007; Herron and Doebeli 2013; Kvitek and Sherlock 2013; Lang *et al.* 2013), genetic drift, admixture and other demographic processes. Excluding clonal interference, which is minimized through our parameter choices in our simulations, none of these other processes are considered in this work, and some of them may also have a substantial qualitative effect on the relationship between future and historical predictability as they may also modify the amount of sign epistasis between the mutations present in an evolutionary path.

The underlying genetic, phenotypic and fitness landscape models used in our simulations are also limited in a number of ways, and could be expanded by including the possibility of multiple adaptive optima, genetically unlinked loci that are capable of adaptation (i.e. recombination between mutations during sexual reproduction), epistasis between loci and the presence of standing genetic variation. Consideration of these processes will likely further complicate the inference of historical predictability. Finally, simulation systems have the advantage of having exact knowledge of the fitness of every genotype. This information must be estimated in natural systems, which may introduce significant noise the inference process. While phenomena such as hidden mutations are likely universal to all of these more complex scenarios, we suspect that the relationship between future and historical predictability may vary between different systems.

Despite our use of a very simple model, we have shown a number of limitations of studying historical predictability when attempting to predict evolution. Not only is historical predictability not directly correlated with future predictability, but it is anticorrelated in our model, suggesting that studying historical predictability may give misleading information about the future predictability of evolution in a given system. In addition, these trends were only discovered through the study of a large number of independent simulations, so the analysis of single adaptive walks is likely of limited utility in systems with highly polygenic adaptation. We also find that the presence of polymorphisms can create historical dependencies in the evolutionary trajectory that are extremely difficult to account for in experimental studies. Thus, this work opens up a number of new questions for study. First, the relationship between future and historical predictability in natural systems is unknown, nor do we know whether this relationship varies between different biological systems. If this relationship does vary, we need to understand the parameters that cause this variation so that one can understand which systems may be amenable to historically predicting evolution and which are not. A similar approach should be taken to understand the conditions under which historical contingency significantly influences adaptive evolution. Our work shows that historical predictability cannot be used as a naive proxy for predicting future evolution, and highlights the need for new approaches to studying future predictability.

## 5. Funding Acknowledgements

The work was funded by the NIH/NHGRI training grant T32 HG000044 and a Stanford Center for Evolutionary and Human Genomics (CEHG) pre-doctoral fellowship to SV, a Stanford Graduate Fellowship and a Stanford CEHG pre-doctoral fellowship to DS and NIH grants RO1 GM115919, GM10036601, GM097415 to DAP. The content of this work is solely the responsibility of the authors and does not necessarily represent the official views of Stanford University or the National Institutes of Health. The authors would like to thank five anonymous reviewers for their valuable feedback while preparing this manuscript.

## Bibliography

Achaz, G., A. Rodriguez-Verdugo, B. S. Gaut, and O. Tenaillon, 2014 The reproducibility of adaptation in the light of experimental evolution with whole genome sequencing. Advances in Experimental Medicine and Biology 781: 211–231.

Altrock, P. M., A. Traulsen, R. G. Reeves, and F. A. Reed, 2010 Using underdominance to bi-stably transform local populations. Journal of Theoretical Biology 267: 62–75.

Blanquart, F., G. Achaz, T. Bataillon, and O. Tenaillon, 2014 Properties of selected mutations and genotypic landscapes under Fisher’s geometric model. Evolution 68: 3537–3554.

Bridgham, J. T., S. M. Carroll, and J. W. Thornton, 2006 Evolution of hormone-receptor complexity by molecular exploitation. Science (New York, N.Y.) 312: 97–101.

Buenrostro, J. D., C. L. Araya, L. M. Chircus, C. J. Layton, H. Y. Chang, et al., 2014 Quantitative analysis of RNA-protein interactions on a massively parallel array reveals biophysical and evolutionary landscapes. Nature biotechnology 32: 562–8.

Crozat, E., C. Winkworth, J. Gaffe, P. F. Hallin, M. A. Riley, et al., 2010 Parallel Genetic and Phenotypic Evolution of DNA Superhelicity in Experimental Populations of Escherichia coli. Molecular Biology and Evolution 27: 2113–2128.

de Visser, J. A. G. M., and J. Krug, 2014 Empirical fitness landscapes and the predictability of evolution. Nature reviews. Genetics 15: 480–90.

Desai, M. M., and D. S. Fisher, 2007 Beneficial mutation selection balance and the effect of linkage on positive selection. Genetics 176: 1759–98.

Fisher, R., 1930 The genetical theory of natural selection. Oxford at the Clarendon Press, Oxford, 1st edition.

Franke, J., A. Klozer, J. A. G. M. de Visser, J. Krug, J. A. G. M. D. Visser, et al., 2011 Evolutionary Accessibility of Mutational Pathways. PLoS Computational Biology 7.

Gould, S. J., 1989 Wonderful Life: The Burgess Shale and the Nature of History. W. W. Norton & Company.

Haldane, J. B. S., 1927 A Mathematical Theory of Natural and Artificial Selection, Part V: Selection and Mutation. Mathematical Proceedings of the Cambridge Philosophical Society 23: 838–44.

Herrera, C. M., C. De Vega, A. Canto, and M. I. Pozo, 2009 Yeasts in floral nectar: A quantitative survey. Annals of Botany 103: 1415–1423.

Herron, M. D., and M. Doebeli, 2013 Parallel Evolutionary Dynamics of Adaptive Diversification in *Escherichia coli*. PLoS Biology 11: e1001490.

Iserbyt, A., J. Bots, H. Van Gossum, and T. N. Sherratt, 2013 Negative frequency-dependent selection or alternative reproductive tactics: maintenance of female polymorphism in natural populations. BMC evolutionary biology 13: 139.

IUCN, 2016 The IUCN Red List of Threatened Species.

Kasumovic, M. M., M. J. Bruce, M. C. B. Andrade, and M. E. Herberstein, 2008 Spatial and temporal demographic variation drives within-season fluctuations in sexual selection. Evolution 62: 2316–25.

Khan, A. I., D. M. Dinh, D. Schneider, R. E. Lenski, and T. F. Cooper, 2011 Negative epistasis between beneficial mutations in an evolving bacterial population. Science 332: 1193–6.

Kimura, M., 1956 Rules for testing stability of a selective polymorphism. Proceedings of the National Academy of Sciences of … 1966: 336–340.

Kimura, M., 1962 On the probability of fixation of mutant genes in a population. Genetics 47: 713–719.

Kimura, M., 1965 A stochastic model concerning the maintenance of genetic variability in quantitative characters. Proc Natl Acad Sci U S A: 731–736.

Kimura, M., and J. F. Crow, 1964 THE NUMBER OF ALLELES THAT CAN BE MAINTAINED IN A FINITE POPULATION. Genetics 49: 725–738.

Kingman, J. F. C., 1961 A mathematical problem in population genetics. Mathematical Proceedings of the Cambridge Philosophical Society 57: 574.

Kojima, K.-i., 1959 Role of epistasis and overdominance in stability of equilibria with selection. Proceedings of the National Academy of Sciences of … 1057: 984–989.

Kvitek, D. J., and G. Sherlock, 2013 Whole Genome, Whole Population Sequencing Reveals That Loss of Signaling Networks Is the Major Adaptive Strategy in a Constant Environment. PLoS Genetics 9: e1003972.

Lang, G. I., D. P. Rice, M. J. Hickman, E. Sodergren, G. M. Weinstock, et al., 2013 Pervasive genetic hitchhiking and clonal interference in forty evolving yeast populations. Nature 500: 571–4.

Lenski, R. E., M. J. Wiser, N. Ribeck, Z. D. Blount, J. R. Nahum, et al., 2015 Sustained fitness gains and variability in fitness trajectories in the long-term evolution experiment with *Escherichia coli*. Proceedings of the Royal Society B: Biological Sciences 282: 20152292.

Levin, B. R. B., J. Antonovics, and H. Sharma, 1988 Frequency-Dependent Selection in Bacterial Populations [and Discussion]. Philosophical Transactions of the Royal Society B: Biological Sciences 319: 459–472.

Lobkovsky, A. E., and E. V. Koonin, 2012 Replaying the tape of life: quantification of the predictability of evolution. Frontiers in Genetics 3: 246.

MacLean, R. C., G. G. Perron, and A. Gardner, 2010 Diminishing returns from beneficial mutations and pervasive epistasis shape the fitness landscape for rifampicin resistance in Pseudomonas aeruginosa. Genetics 186: 1345–54.

Malcolm, B., K. Wilson, and B. Matthews, 1990 Ancestral lysozymes reconstructed, neutrality tested, and thermostability linked to hydrocarbon packing. Nature 345: 86–89.

Mandel, S. P. H., 1959 THE STABILITY OF A MULTIPLE ALLELIC SYSTEM. Heredity 13: 289–302.

Martin, G., 2014 Fisher’s Geometrical Model Emerges as a Property of Complex Integrated Phenotypic Networks. Genetics 197: 237–255.

Martin, G., S. F. Elena, and T. Lenormand, 2007 Distributions of epistasis in microbes fit predictions from a fitness landscape model. Nature genetics 39: 555–60.

Orr, H. A., 1999 The evolutionary genetics of adaptation: a simulation study. Genetical Research 74: 207–14.

Orr, H. A., 2005 The genetic theory of adaptation: a brief history. Nature reviews. Genetics 6: 119–27.

Ortlund, E. A., J. T. Bridgham, M. R. Redinbo, and J. W. Thornton, 2007 Crystal structure of an ancient protein: evolution by conformational epistasis. Science (New York, N.Y.) 317: 1544–8.

Palmer, M. E. M., A. Moudgil, and M. W. Feldman, 2013 Long-term evolution is surprisingly predictable in lattice proteins. Journal of the Royal Society, Interface 10: 20130026.

Rainey, P., and M. Travisano, 1998 Adaptive radiation in a heterogeneous environment. Nature 394: 69–72.

Roy, S. W., 2009 Probing evolutionary repeatability: neutral and double changes and the predictability of evolutionary adaptation. PLoS ONE 4: e4500.

Saltz, J. B., and S. V. Nuzhdin, 2014 Genetic variation in niche construction: implications for development and evolutionary genetics. Trends in ecology & evolution 29: 8–14.

Sellis, D., B. B. J. Callahan, D. A. Petrov, and P. W. Messer, 2011 Heterozygote advantage as a natural consequence of adaptation in diploids. Proceedings of the National Academy of Sciences of the United States of America 108: 20666–71.

Sousa, A., S. Magalhaes, and I. Gordo, 2012 Cost of antibiotic resistance and the geometry of adaptation. Molecular Biology and Evolution 29: 1417–1428.

Tenaillon, O., 2014 The Utility of Fisher’s Geometric Model in Evolutionary Genetics. Annual Review of Ecology, Evolution, and Systematics 45: 179–201.

Tenaillon, O., J. E. Barrick, N. Ribeck, D. E. Deatherage, J. L. Blanchard, et al., 2016 Tempo and mode of genome evolution in a 50,000 generation experiment. Nature 536: 156–170.

Tenaillon, O., A. Rodriguez-Verdugo, R. L. Gaut, P. McDonald, A. F. Bennett, et al., 2012 The Molecular Diversity of Adaptive Convergence. Science 335: 457–461.

Tenesa, A., P. Navarro, B. J. Hayes, D. L. Duffy, G. M. Clarke, et al., 2007 Recent human effective population size estimated from linkage disequilibrium. Genome Research 2: 520–526.

Venkataram, S., B. Dunn, Y. Li, A. Agarwala, J. Chang, et al., 2016 Development of a Comprehensive Genotype-to- Fitness Map of Adaptation-Driving Mutations in Yeast. Cell: 1–12.

Weinreich, D. M., N. F. Delaney, M. A. Depristo, and D. L. Hartl, 2006 Darwinian evolution can follow only very few mutational paths to fitter proteins. Science 312: 111–4.

Weinreich, D. M., R. A. Watson, and L. Chao, 2005 Perspective:Sign Epistasis and Genetic Constraint on Evolutionary Trajectories. Evolution 59: 1165.

Wiser, M. J., N. Ribeck, and R. E. Lenski, 2013 Long-Term Dynamics of Adaptation in Asexual Populations. Science (New York, N.Y.) 1364.

Wright, S., 1931 Evolution in Mendelian Populations. Genetics 16: 97–159.

